# Towards standards for analysing structural variants from short-read whole genome sequencing data

**DOI:** 10.64898/2026.10.13.758836

**Authors:** James C. Slimmer, Johanna L. Jones, William R. Reay, Kathryn P. Burdon, Nicholas B. Blackburn

**Affiliations:** Menzies Institute for Medical Research, University of Tasmania, Hobart, TAS 7000, Australia

## Abstract

No widely accepted standard exists for analysing structural variants from short-read whole genome sequencing (WGS) data. We use comprehensive SV truth sets (HG002 T2T-Q100, Platinum Pedigree) and long-read sequencing databases (1KGP-ONT, CoLoRS) to benchmark SV breakpoints and sizes at base pair resolution. Using 30X Illumina WGS data, only five of sixteen individual tools tested reliably output deletions, insertions and duplications that perfectly match the truth sets. Ensemble pipelines (e.g. GATK-SV, MOPline, Parliament2), frequently used in population-scale datasets, reduced perfect matching accuracy by up to 98% compared to their highest-performing constituent tools. This inaccuracy propagates into a short-read database (gnomAD-SV) constructed from an ensemble pipeline (GATK-SV) and likely affects other databases (TOPMed, All of Us, Biobank Japan). As a proof-of-concept, we demonstrate an intra-sample merging strategy that generates the highest F1-scores for deletions and insertions on the HG002 sample, recovering more perfect matches than total matches from the best performing ensemble pipeline tested. Common HG002 truth SVs (allele frequency > 1%) showed extremely high concordance for exact matches between our ensemble callset and two orthogonal long-read sequencing databases (CoLoRS, ONT-1KGP). This study provides a vital step towards robust standards for SV analysis, guiding short-read WGS ensemble pipeline development and improving cross-study comparisons.

## Introduction

Structural variants (SV) are defined as DNA sequence rearrangements of 50 base pairs or greater, encompassing insertions, deletions, duplications, inversions and translocations. Evidence for the contribution of SVs to both monogenic diseases and complex traits has grown significantly in recent years [1]. Long-read whole genome sequencing (WGS) is now also widely recognised as necessary for comprehensive SV discovery [2]. However, due to cost and technical factors, such as sample requirements, the workhorse of modern genomics for analysing all types of genetic variation will continue to be short-read WGS for the foreseeable future [3–6].

A plethora of short-read SV calling tools and pipelines have been developed over the past fifteen years [7–10]. Unlike single nucleotide variants (SNV) and small indels less than 50 base pairs [11,12], there is no gold standard SV calling tool or pipeline, nor is there a consistent standard for benchmarking performance or comparing SVs between callsets. Accurate discovery and comparison of SVs is inherently challenging due to read length constraints and the diverse types of read evidence (split-read, discordant paired-end, and read depth) utilised by different algorithms [13]. For example, tools relying heavily or solely on read depth are expected to produce imprecise SV breakpoints. To facilitate “fair” comparisons between different tools, SV benchmarking studies have traditionally used highly permissive matching thresholds. Consequently, varying definitions of when SVs should be considered the same underlying variant has hindered the development of robust benchmarking standards.

Despite these challenges, previous benchmarking studies consistently show that the performance of individual callers varies by SV type and size, particularly for insertions [14,15]. To maximise detection sensitivity, ensemble pipelines integrating multiple tools have been widely adopted by population-scale and cohort sequencing projects [5,9,10,16–20]. However, this increased sensitivity comes at the cost of methodological complexity, requiring careful selection of individual tools to balance sensitivity while controlling for an inflation of false positives. Furthermore, some existing pipelines are proprietary [7], lack regular updates [8], or are primarily designed for cloud platforms [9], all of which are barriers to broader uptake by the genomics community. As a result, many studies have developed project-specific, *ad-hoc* pipelines with custom quality control, SV discovery, merging and genotyping [16,17,19,20]. The proliferation of custom ensemble pipelines compounds problems related to the lack of robust benchmarking standards. Determining how to accurately merge redundant SV representations output from multiple tools also requires defining when two or more SVs represent the same underlying variant (intra-sample merging) and predicting which representation is correct. We define ‘representation’ here as the true, non-redundant exact breakpoint and size of an SV within a sample relative to the reference genome used.

Ultimately, best practices in SV analysis should consider not only the comprehensiveness of SV discovery, but also the accuracy of the breakpoints and sizes. Under the current permissive matching paradigm, the accuracy of SV representations in most short-read studies is unclear. Applying a single permissive matching threshold obscures performance at base pair resolution and potentially masks true genetic variation across samples and studies. By altering matching parameters and thresholds, population studies can report greater diversity or overlap of SVs, complicating cross-study comparisons. Exact breakpoints and sizes are critical for SV interpretation: they determine whether an SV has exonic impacts, causes a frameshift, or disrupts a splice site or regulatory element. Imprecise breakpoints compromise SV interpretation; for instance, a rare, pathogenic frameshift deletion could be falsely disregarded as a common, benign in-frame deletion if they are assumed to be the same SV simply because they differ by only one base pair.

The growing availability of comprehensive SV truth sets and long-read sequencing population databases provides opportunities for analysing SVs at base pair resolution with short-read WGS data. To this end, we benchmarked sixteen SV callers and four ensemble pipelines against two recently developed SV truth sets (HG002 T2T-Q100 [21,22]; Platinum Pedigree [23]) using matched short-read sequencing data and increasingly stringent matching thresholds. We further analysed the breakpoint and size accuracy of short-read WGS-derived HG002 SV representations against orthogonal long-read (1KGP-ONT; CoLoRS) and short-read (gnomAD-SV) population databases, evaluating both callset-specific comprehensiveness and the concordance of common (allele frequency > 1%) representations intersected across callsets and databases.

## Materials and Methods

### Structural variant truth VCF preprocessing

The Platinum Pedigree SV truth set VCF (with tandem repeats included) was obtained from s3://platinum-pedigree-data/variants/merged_sv_truthset/GRCh38/. The NA12877 and NA12878 samples from the GRCh38 Platinum Pedigree SV truth set VCF were split into separate VCFs with homozygous reference genotypes excluded using bcftools view -s {sample} -e ‘FORMAT/GT=”0/0”’. Upon manual inspection of the truth VCFs, we found duplicated and unmerged SVs at some loci that appeared to coincide with the presence of multiple Ns in the reference or alternate sequences. These variants were removed except when the only reference base was a single N, which represents a Sniffles2 call. There were 23,828 and 23,809 SVs remaining in the truth set VCFs for NA12878 and NA12877, respectively, after filtering.

The GRCh38 HG002 T2T-Q100 v0.020 SV truth VCF was obtained from https://ftp-trace.ncbi.nlm.nih.gov/ReferenceSamples/giab/data/AshkenazimTrio/analysis/NIST_HG002_DraftBenchmark_defrabbV0.020-20250117/. Insertions and deletions greater than 49bp and ALT alleles containing “*” were removed from the HG002 T2T-Q100 SV truth VCF using bcftools. Any SV record with N in the REF or ALT sequences was also removed for consistency with our Platinum Pedigree truth set filtering. After processing, there were 28,174 SVs within the high confidence regions of the HG002 truth set VCF.

### Structural variant callsets

Downsampled PCR-free 151bp paired-end Illumina WGS FASTQ files for HG002 were downloaded from https://console.cloud.google.com/storage/browser/brain-genomics-public/research/sequencing/fastq/novaseq/wgs_pcr_free/30x. The PCR-free 150bp paired-end Illumina WGS FASTQ files for NA12878 and NA12877 were downloaded from https://www.ebi.ac.uk/ena/browser/view/PRJEB36890 and https://www.ebi.ac.uk/ena/browser/view/PRJEB31736, respectively. Read alignment (BWA-MEM v0.7.18 and hg38 with alt contigs) and deduplication was conducted using the nf-core/sarek pipeline (v3.4.4) with default parameters, except --skip baserecalibrator was included to turn base recalibration off. Sequencing reads were also deduplicated using both fastp --disable_quality_filtering --dedup_accuracy 6 and Picard MarkDuplicates. The proportion of total bases required to reach 30X coverage was calculated from sarek’s mosdepth output, which uses the deduplicated reads mapped by BWA-MEM as input. The 30X downsampling value was input to the command seqtk sample -s100 {fastq} {downsample_proportion}.

To simplify installation and facilitate SV calling and benchmarking, callsets for some tools were acquired from the Parliament2 and MOPline ensemble caller outputs. A list of all callers, versions and pipeline-specific SV caller outputs used for benchmarking can be found in Supplementary Table 1. All structural variant callers were used in the Parliament2 pipeline except Breakseq2 due to a fault preventing output of SV calls. Additionally, the size of every SV in the Parliament2 callsets was rounded to the nearest integer. GATK-SV callsets for NA12878 and NA12877, and HG002 were obtained from https://ftp.1000genomes.ebi.ac.uk/vol1/ftp/data_collections/1000G_2504_high_coverage/working/20210124.SV_Illumina_Integration/ and https://github.com/broadinstitute/blend_seq_paper, respectively. The NA12878 and NA12877 samples from the high coverage whole genome resequenced 1KGP GATK-SV VCFs were extracted with homozygous reference genotypes excluded using bcftools view -s {sample} -e ‘FORMAT/GT=”0/0”’ -g ^miss. CNV type SVs were also removed from the GATK-SV VCFs. Manual inspection of calls in the INSurVeyor VCFs revealed a systematic off-by-one coordinate error compared to the truth set insertions. All INSurVeyor calls were therefore modified by subtracting 1 from the VCF END field. Each MOPline callset was generated according to developer instructions from https://github.com/stat-lab/MOPline with the optional filter step also applied to the final output VCFs. The MOPline11t callset includes all SV callers supported by preset conversion scripts — MOPline6t refers to the 6tools_2 preset.

### Structural variant benchmarking

Truvari v5.3.0 [24] was used for all SV benchmarking, merging and database comparisons. There are many methods for increasing precision of SV callsets. Since we were interested in accurate matches, we used all SVs from the main VCF output by each caller or pipeline where possible (Supplementary Table 1). We applied custom scripts utilising the pysam v0.22.1 library (**Data Availability**) to filter and format SV caller outputs into VCFs compatible with Truvari. Briefly, VCF headers were added or modified to standardise formats across callsets. Because Truvari bench prioritises using END coordinates over INFO/SVLEN, and some callsets had either missing END coordinates or values that did not conform to VCF 4.2 specifications, every END coordinate for deletions and duplications was recalculated as the absolute value of SVLEN + POS. If a deletion or duplication did not have a length, then it was calculated as END - POS when an END coordinate was present. Deletions and insertions/duplications were benchmarked separately by splitting truth and query VCFs with bcftools view -i ‘INFO/SVTYPE’. The following Truvari bench commands were used for deletions and insertions/duplications:

Truvari bench -s 50 -S 50 --pick single --sizemax 10000000

The HG002 T2T-Q100 high confidence BED regions (GRCh38_HG2-T2TQ100-V1.1_stvar.benchmark.bed) were used with --includebed when benchmarking HG002 callsets. The --dup-to-ins option was used to convert duplications into point insertions since duplications are represented as sequence-resolved insertions in the Platinum Pedigree and HG002 T2T-Q100 truth VCFs. Only a single match (--pick single) was allowed between any detected SV or truth set SV. We set --sizemax to 100000 when benchmarking SVs with the NA12878 and NA12877 samples because the Platinum Pedigree truth VCFs do not contain SVs > 100kb.

### Merging structural variant callsets

To construct the priority_merge callset, individual caller VCFs were first split by SV type (insertions/duplications and deletions) and partitioned into size bins using bcftools view. For deletions, callers were prioritised by size bin in the following descending order: <500bp (gcSV, Manta, SurVIndel2), 500-1000bp (gcSV, SurVIndel2, Manta), and ≥1kb (SurVIndel2, Manta, gcSV). SurVIndel2 was chosen over DELLY as the third caller because it demonstrated superior recall in the 50-200bp range, where most non-redundant deletions are found. For insertions, the size bins and descending priority order was: <200bp (gcSV, Manta, INSurVeyor), 200-1000bp (gcSV, INSurVeyor, Manta), and ≥1kb (INSurVeyor, gcSV, Manta).

Intra-sample merging was conducted by sequentially combining caller VCFs based on the priority orders above. Each caller VCF was split by SV type (DEL for deletions, DUP/INS for insertions) and size using bcftools view. For every SV type and size bin, the VCF from the highest-priority caller was used as the initial base VCF. Next, the VCF from the second-priority caller was compared against the base VCF with Truvari bench using the previously described benchmarking parameters and the most permissive matching threshold (70% reciprocal overlap for deletions; 500bp reference distance and 70% size for insertions). SVs from the second caller that did not match the base VCF — output by Truvari as false positives (fp.vcf.gz) — were extracted and appended to the base VCF using bcftools concat to create an updated, non-redundant base VCF. This process was then repeated for the third-priority caller against the updated base VCF. Finally, the fully merged, non-redundant VCFs across all size bins were combined (bcftools concat) and sorted (bcftools sort) to produce the priority_merge analysis VCF for each SV type.

### Repeat element annotations

SVs in the HG002 T2T-Q100 truth VCF were annotated using the SimpleRepeats and SegmentalDups tracks from the UCSC Genome Browser [25]. The track files (simpleRepeat.txt.gz and genomicSuperDups.txt.gz) were downloaded from the UCSC hg38 database (https://hgdownload.soe.ucsc.edu/goldenPath/hg38/database/). Bedtools merge was first used to flatten overlapping coordinates within the respective track BED files. Bedtools subtract was then applied before concatenating the files to produce exclusive annotation coordinates. If any part of a deletion overlapped with an annotated coordinate, or if an insertions reference coordinate fell within one of these regions, the SV was labelled as Repeat/SegDup. All remaining SVs were labelled as Unique.

### Matching structural variants to 1KGP-ONT, gnomAD-SV and CoLoRS databases

The CoLoRS v1.2, gnomAD-SV v4.1 non-neuro controls and 1KGP-ONT VCFs were obtained from https://zenodo.org/records/14814308, https://gnomad.broadinstitute.org/downloads, and https://s3.amazonaws.com/1000g-ont/needLR/, respectively. One base pair was added to the END coordinate of deletions in the 1KGP-ONT VCF to correct an off-by-one file format error between the SVLEN and POS values. The END coordinate for insertions in the gnomAD-SV VCF was also modified to match the POS coordinate as per VCF v4.2 specifications.

The true positive base VCFs output by Truvari were intersected to identify the subset of HG002 truth set SVs that were present in all three callsets (Manta, GATK-SV, priority_merge) at permissive matching thresholds (deletions: 70% reciprocal overlap; insertions/duplications: 500bp reference distance, 70% size). The Truvari MatchIds from the respective true positive comparison VCF were then used to extract the original VCF records, followed by matching these calls to gnomAD-SV, 1KGP-ONT and CoLoRS SV database VCFs at permissive matching thresholds as well. The GATK-SV, Manta and priority_merge callset representations (genomic coordinates and sizes only) were then intersected with their database SV matches. To be considered for the downstream analysis of common SVs, we required two conditions to be met: 1) the original Manta, GATK-SV and priority_merge callset representations must all match the same SV record in each database, and 2) the allele frequency of the matched SV must be 1% or greater in each database. To evaluate missingness and matching accuracy against the population databases, we matched all Manta, GATK-SV and priority_merge callset representations of the HG002 truth SVs detected. The callset representations were annotated for sequence context (Repeat/SegDup vs. Unique) by mapping them back to the UCSC SegmentalDups and SimpleRepeats track annotations of the underlying HG002 truth SVs. Perfect matches were defined by values of 0 in the Truvari SizeDiff, StartDistance and EndDistance VCF INFO fields.

## Results

### Ensemble pipelines output fewer perfectly matched SVs than individual callers

Publicly available Illumina PCR-free FASTQ files for three samples (HG002, NA12878, and NA12877) were downsampled to 30X coverage and processed with the nf-core sarek pipeline [26]. Sixteen SV calling tools and three ensemble pipelines were then applied to the deduplicated CRAM files. Callsets from the GATK-SV ensemble pipeline generated by previous studies for HG002 [27], and NA12878 and NA12877 [5] were also included (**Methods**). Unless stated otherwise, values related to Figures 1 and 2 indicate averages (mean) across the three sample truth sets. First, we examined the matching accuracy of deletions using Truvari bench with increasingly stringent reciprocal overlap thresholds (70-100%) between the truth set and the deletions output by a specific caller or pipeline (**Figure 1A**). In each sample, Manta, gcSV, SurVIndel2 and DELLY detected over 20% of the truth set deletions with perfect reciprocal overlap. The fifth best individual caller (GRIDSS2) for perfect deletion matches only achieved a maximum recall of 4.0%. Seven of the fourteen callers output deletions where less than 1% of their true positives were exact matches to the sample truth set. Notably, the Parliament2, GATK-SV, MOPline6t and MOPline11t ensemble pipelines (which all include Manta) produced substantially fewer deletion matches at 100% reciprocal overlap compared to Manta individually. On average across the three samples, Parliament2 output 73.7% less deletions than Manta as perfect matches (Manta: 2970, Parliament2: 781), whereas GATK-SV, MOPline6t and MOPline11t did not generate more than 6.5% as many as Manta in any sample. Moreover, except for Parliament2, and despite ensemble pipelines including multiple tools for detecting deletions, recall for Manta at 100% reciprocal overlap was higher than GATK-SV, MOPline6t and MOPline11t in every sample when compared to the lowest matching threshold (70% reciprocal overlap) we tested. Manta recall at 70% reciprocal overlap was still remarkably similar to Parliament2. Additionally, gcSV detected more perfectly matched deletions than the deletions output by Parliament2 at the lowest matching threshold; the NA12878 and NA12877 samples had comparable recall between these thresholds. We also surprisingly found MOPline6t recall was higher than MOPline11t at every matching threshold, even though three additional tools that detect deletions were used in MOPline11t to generate this callset.

**Figure 1.**
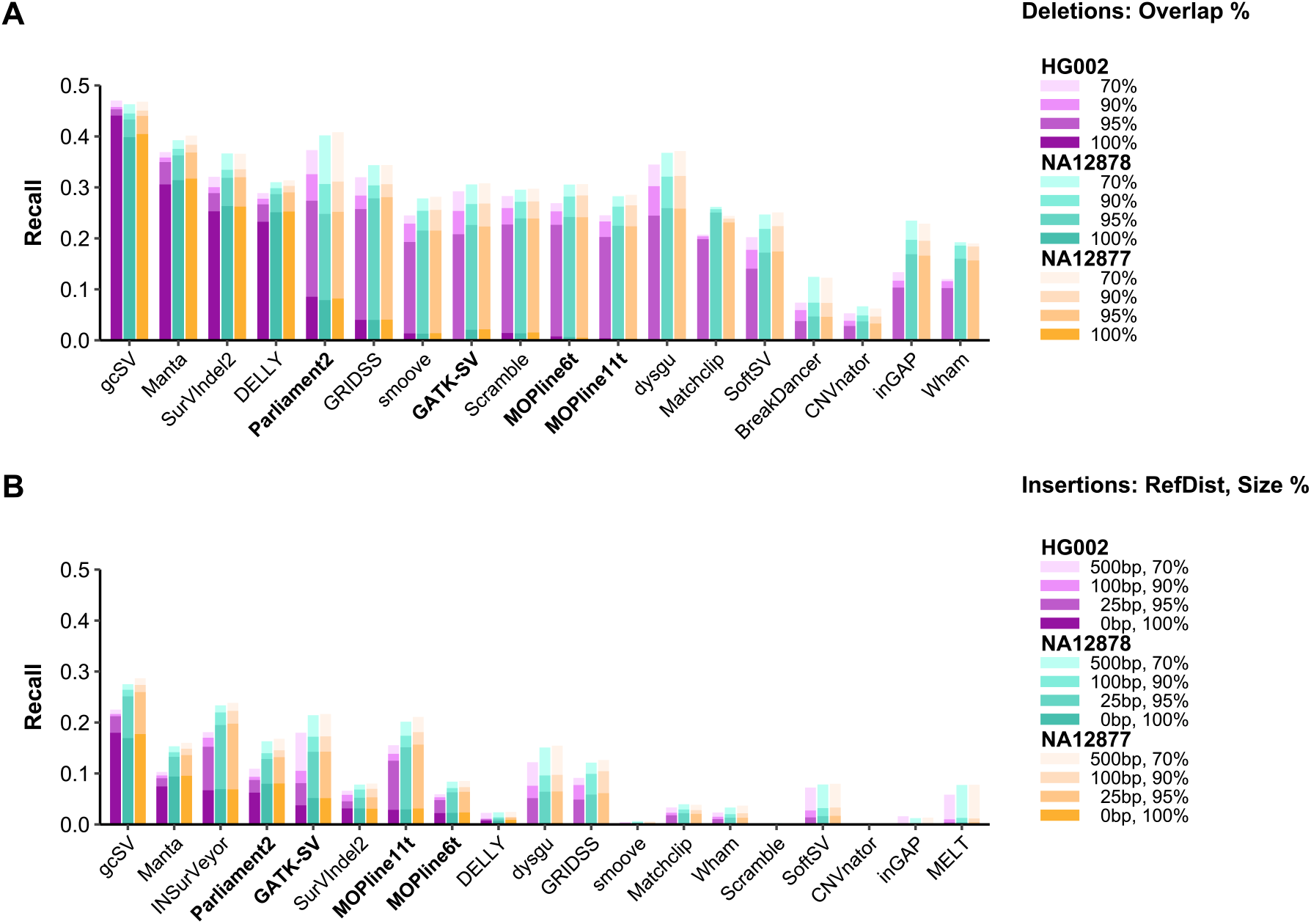
Recall of insertions and deletions at different matching thresholds. Each sample is indicated by a different colour. From lightest to darkest, the shade of each colour represents increasingly stringent matching thresholds. **A)** Recall of deletion calls ordered by descending mean F1-score (harmonic mean of recall and precision) at 100% reciprocal overlap. **B)** Recall of insertion/duplication calls ordered by descending mean F1-score for exact reference coordinate and size matches. Ensemble pipelines that integrate multiple SV calling tools are highlighted in bold (all include Manta).

**Figure 2.**
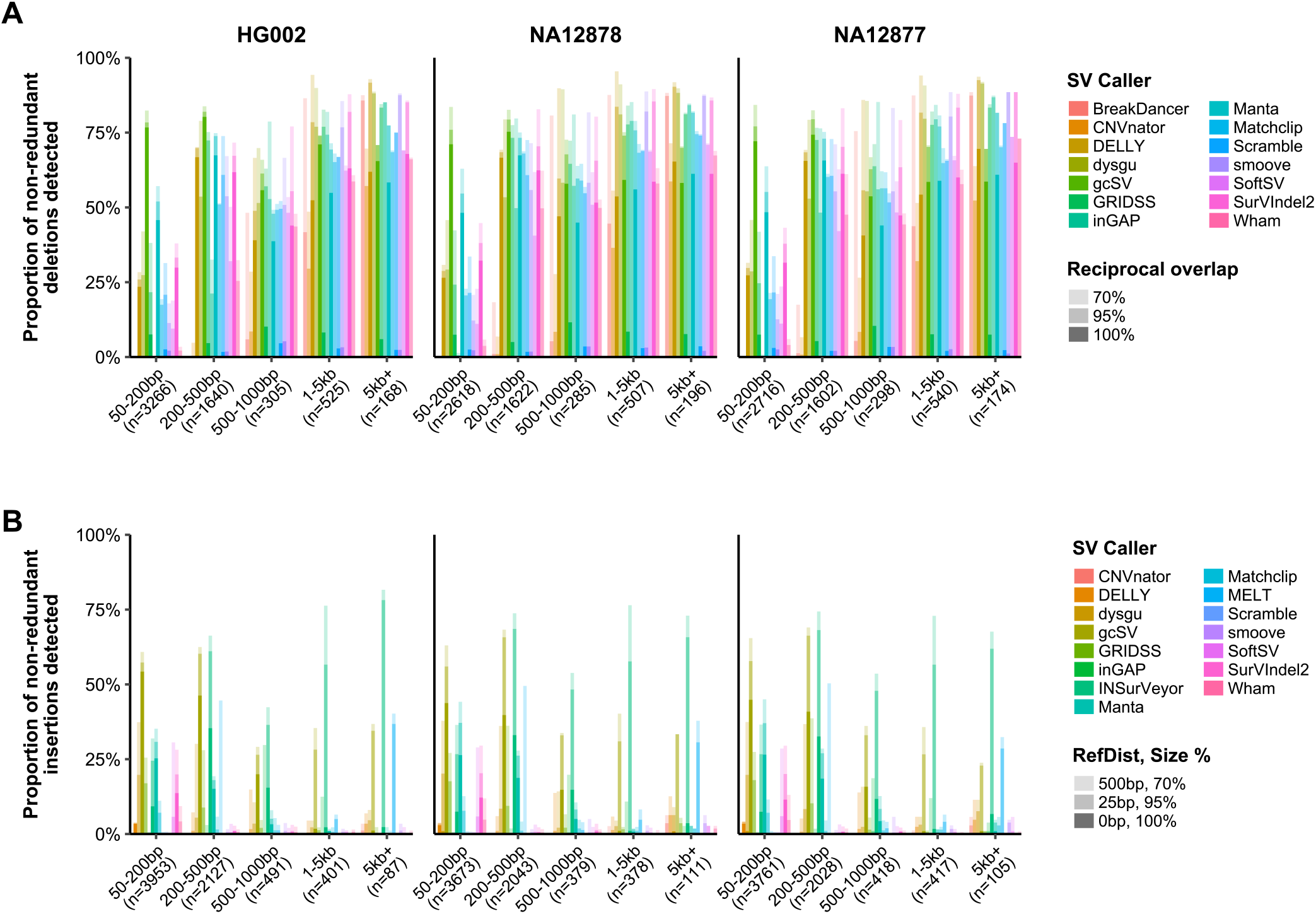
Size stratification of deletions and insertions by matching threshold. From lightest to darkest, the shade of each colour represents increasingly stringent matching thresholds. For each size bin, the proportion of non-redundant SVs detected by a caller is calculated as the number of truth set SVs detected by that caller divided by the total number of non-redundant truth set SVs detected across all callers in that bin. **A)** All non-redundant deletions detected in each sample truth set binned by size. **B)** All non-redundant insertions/duplications detected in each sample truth set binned by size.

Next, we assessed insertions and duplications using breakpoint distance from the reference insertion point and insertion size as matching parameters. Duplications were converted to point insertions for consistency with their representation in the truth set VCFs (Methods). Unlike deletions, no tool or pipeline could perfectly match at least 20% of the insertions in any sample’s truth set; in fact, only one out of nineteen (gcSV) produced more than 20% of insertions at the most permissive matching threshold (±500bp reference distance, 70% size) in all three sample callsets (Figure 1B). The ensemble pipelines generally outperform Manta in recall for insertions at less stringent matching thresholds. Yet, when comparing perfect matches of individual tools to ensemble pipelines, we observe the same phenomenon previously seen with deletions. Parliament2 produced 15.7% less perfect matches than Manta individually (Manta: 1369, Parliament2: 1154). This discrepancy was much larger for the GATK-SV (-46.9%), MOPline11t (-65.8%) and MOPline6t (-73.9%) callsets. The smaller discrepancy between Parliament2 and other ensemble pipelines can be explained by Parliament2 using callers that output relatively fewer duplications and insertions that are redundant with the Manta calls. For example, MOPline11t integrates five more SV callers than MOPline6t (both include Manta) and detects 248.8% more insertions overall. The increase in insertions detected by MOPline11t is likely driven by the inclusion of INSurVeyor, considering the greater individual performance of this tool. However, when comparing recall at the perfect match threshold, MOPline11t (2.97% total) only increased by 31.1% compared to MOPline6t (2.27% total), whereas recall at the same threshold for the individual Manta and INSurVeyor callsets was 8.79% and 6.83%, respectively. Similar to Parliament2, the increase in perfectly matched insertions between MOPline11t and MOPline6t could be attributable to unique insertions that were only detected by INSurVeyor.

### The best performing caller varies by SV type, size and matching threshold

Collectively, the results from insertions and deletions suggest a fundamental problem with the implementation of intra-sample merging strategies in ensemble pipelines – namely, the selection of representative breakpoints and sizes from redundant SVs identified between multiple callers. Previous benchmarking studies show recall varies across SV types and size ranges [14,15]. Given the breakpoint and size inaccuracy observed for some individual tools included in ensemble pipelines (Figure 1), we stratified all non-redundant insertions and deletions detected in the sample truth sets by size, type and matching threshold. Non-redundant SVs are the truth set SVs detected by at least one caller (i.e. uniquely detected truth set SVs). Non-redundant SV proportions were calculated per bin using the total number of unique SVs detected in the sample truth set as the denominator (Figure 2). As expected, we found specific callers perform better than others depending on SV type, size and matching threshold. Across all tools tested, there were 5904, 5228, and 5330 non-redundant deletions detected in the HG002, NA12878 and NA12877 truth sets at the 70% reciprocal overlap threshold, respectively (Figure 2A). In the 50-500bp range, gcSV and Manta consistently detected the highest proportion of non-redundant deletions at the perfect matching threshold in each sample truth set. There is a substantial difference in the proportion of perfectly matched non-redundant deletions between gcSV and Manta in the 50-200bp range (gcSV: 77.1%; Manta: 47.4%). This is less pronounced between 200-500bp, but gcSV still found 14.7% more perfect matches than Manta. From 500bp-1kb, gcSV output the most perfectly matched deletions (55.8%); however, SurVIndel2 outperforms Manta in this size range (SurVIndel2: 47.6%; Manta: 42.5%). Our capability to infer the best performing callers for perfect deletion matches is less clear above 1kb.

There were 7059, 6584, and 6729 non-redundant insertions detected across all tools in the HG002, NA12878 and NA12877 truth sets, respectively, that matched within 500bp of their reference insertion point at 70% size similarity (Figure 2B). In both the 50-200bp and 200-500bp, gcSV output the most insertions perfectly matching the truth set in every sample (minimum 39.7%). Manta detected the second highest proportion of non-redundant insertions for 50-200bp (gcSV: 47.6%; Manta: 26.0%), whereas INSurVeyor generated more perfect matches than Manta between 200-500bp (INSurVeyor: 33.7%; Manta: 17.4%). In two out of three samples, gcSV also detects the largest proportion of perfectly matched insertions in the 500bp-1kb size range (HG002: 20.0%; NA12877: 15.8%) and had proportions equal to the second best caller (INSurVeyor: 14.8%) for NA12878. Consistent with the limitations of short sequencing read lengths [13], matching accuracy for insertions dramatically decreases above 1kb. The proportion of perfectly matched insertions detected by the best caller drops from 16.8% at 500bp-1kb (gcSV), to 1.66% at 1-5kb (INSurVeyor) and 4.19% at 5kb+ (INSurVeyor). Despite this, INSurVeyor detects a vastly higher proportion of non-redundant insertions above 1kb at the second highest match threshold (95% size; ±25bp reference distance) compared to the next best caller (gcSV). In the 1-5kb range, 28.6% of insertions were detected by gcSV whereas 57.0% were detected by INSurVeyor. For insertions above 5kb, 30.2% and 68.6% were detected by gcSV and INSurVeyor, respectively.

### A benchmarking-informed caller priority order increases intra-sample merging accuracy

In practice, when using multiple callers and conducting intra-sample merging, we do not know what the ‘true’ representation is between the two or more SVs that have been matched together. One of the simplest methods is to use the benchmarking performance of each caller within SV type, size and matching threshold stratifications to establish a priority order for which caller is more likely to have the most accurate representation of the SV between the redundant matches. Since, to our knowledge, this type of functionality is not readily implemented in any SV merging tool, we adopted a crude merging approach to generate a ‘priority_merge’ callset by splitting individual caller VCFs by SV type and size, followed by sequentially matching and combining true and false positive output VCFs from Truvari bench as a proof of concept (Methods). The priority orders used can be found in the Methods section. We then compared our ensemble callsets (priority_merge) for each sample to the individual tools used and the four ensemble pipelines from Figure 1 by benchmarking them against their sample truth sets (Figure 3, Supplementary Figures 1 & 2). The HG002 priority_merge callset had the highest F1-scores and recall at all matching thresholds for both insertions (gcSV + Manta + INSurVeyor) and deletions (gcSV + Manta + SurVIndel2), despite the decrease in precision that occurs by including all false positives from multiple callers (Figure 3A & Figure 3B). Additionally, except for insertions in the gcSV callset, deletion and insertion recall for priority_merge at the perfect match threshold was higher than the recall of any callset at the lowest match threshold. Out of 5904 non-redundant deletions that were detected across all callers in the HG002 sample truth set, 82.2% were present in the priority_merge callset as perfect matches and 92.8% matched at the 70% reciprocal overlap threshold. For non-redundant insertions, 50.3% (3549/7059) were found as perfect matches in the HG002 priority_merge callset and 73.1% (5158/7059) were detected at the 500bp reference distance and 70% size threshold. Similar improvements in recall, F1-scores, and non-redundant insertion and deletion detection across matching thresholds were observed in the priority_merge callsets for NA12878 and NA12877 (Supplementary Figure 1 & 2). However, the NA12878 and NA12877 samples had lower precision and less recall improvement at the perfect match threshold relative to HG002, which resulted in marginally lower F1-scores than gcSV. Nevertheless, for deletions and insertions, the NA12878 and NA12877 priority_merge ensemble callsets had the highest recall and F1-scores at the second most stringent matching threshold while increasing the recall of perfect matches above their individual callers. This indicates comprehensive single sample callsets can be achieved in general using this approach without sacrificing matching accuracy. The ‘priority_merge’ callsets we generated also had higher or comparable F1-scores and precision to the other tools and pipelines without applying any method to reduce false positives.

**Figure 3.**
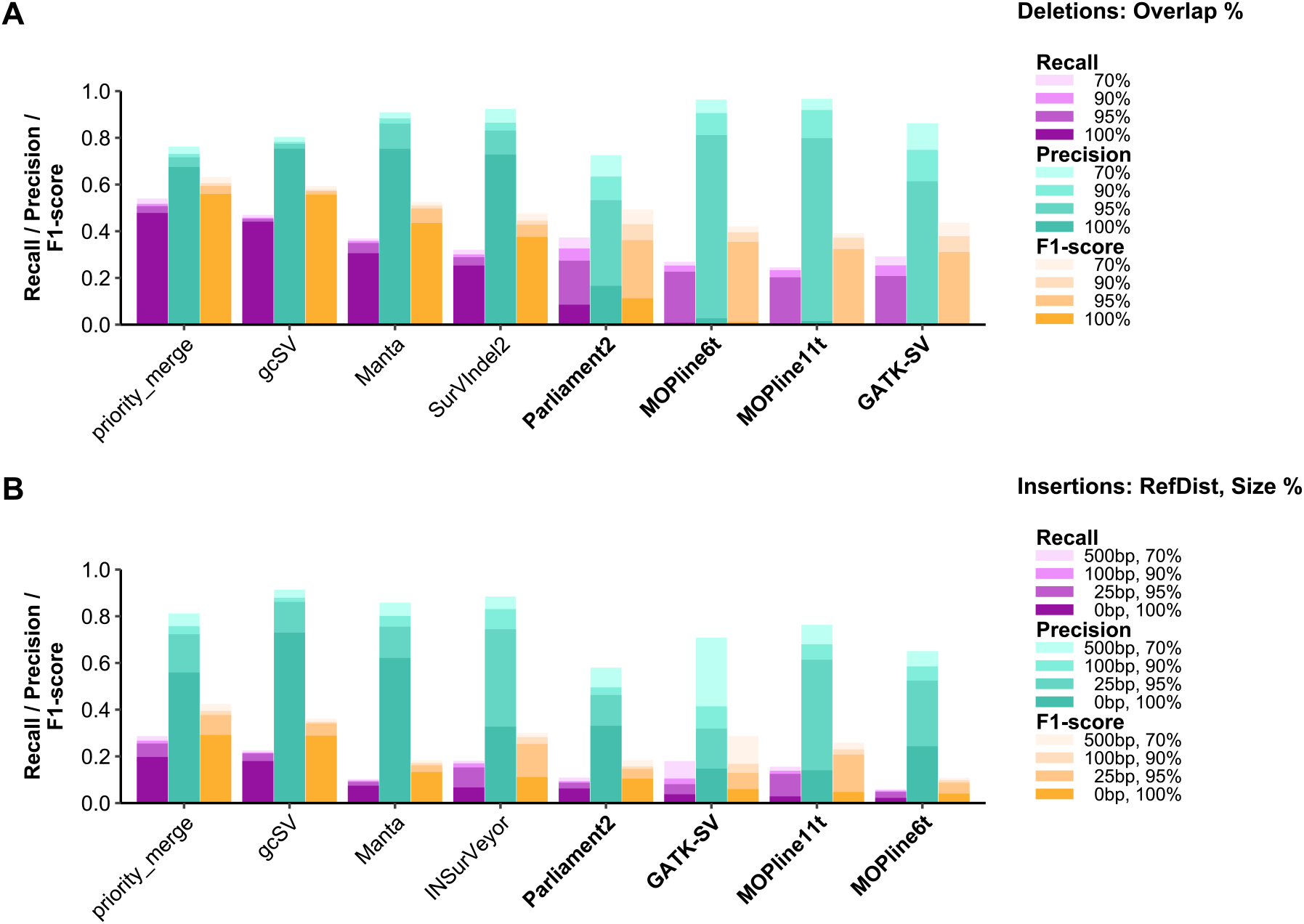
Selecting a representation from redundant structural variant matches using a benchmarking-informed caller priority order increases matching accuracy. **A** ‘priority_merge’ callset was constructed for the HG002 sample by splitting individual caller VCFs by SV type and size bin, followed by merging them with Truvari using a caller priority order (e.g. 50-500bp deletions: gcSV > Manta > SurVIndel2; 50-200bp insertions: gcSV, Manta, INSurVeyor) informed by their benchmarking performance. The caller priority orders for the priority_merge callsets can be found in the **Methods** section. HG002 callsets split into **A)** deletions and **B)** insertions are ordered by descending F1-score at the perfect match threshold. Colours indicate the recall, F1-score and precision for each callset along the x-axis. From lightest to darkest, the shade of each colour represents increasingly stringent matching thresholds. Ensemble pipelines that integrate multiple SV calling tools are highlighted in bold (all include Manta).

### Ensemble pipeline breakpoint and size inaccuracies propagate into an SV database

To investigate the matching accuracy of ensemble methods relative to an individual caller further, we examined common insertions and deletions in two long-read sequencing databases (1KGP-ONT – 500 high coverage nanopore resequenced 1KGP samples [28]; CoLoRS - 1,427 PacBio HiFi sequenced samples from the Consortium of Long Read Sequencing) and a short-read sequencing database (gnomAD-SV). We first intersected the HG002 truth SVs detected in three callsets (Manta, GATK-SV and priority_merge) and extracted the callset-specific representations of their shared truth SVs (Methods). The three callset representations for each of the shared HG002 truth SV were then matched to the three SV databases. If the same three callset representations were found to match as a set to a single SV in each of the three databases, then they were retained for further analysis. Additionally, we further restricted the SVs analysed by requiring the database-matched SVs to have an allele frequency (AF) > 1% in all three databases. Altogether, this generated an analysis set of SVs that are observed in the HG002 truth set, present in the 1KGP-ONT, CoLoRS and gnomAD-SV databases (with varying representations), and occurring at a frequency in each database >1%. To stratify SVs by sequence context, we annotated the HG002 truth SVs with SimpleRepeats and SegmentalDups UCSC track information [25] prior to the matching and intersections. Matching between callsets and databases used the most permissive matching threshold from Figures 1-3 unless stated otherwise. The Manta and priority_merge callsets output perfect matches to less than 25% (n=1,874) of common (AF > 1%) HG002 deletions found in gnomAD-SV (Figure 4A). The number of perfect matches from Manta and priority_merge to gnomAD-SV roughly doubled for deletions in genomic regions annotated as Unique (not in SimpleRepeats or SegmentalDups). It should be noted that Unique does not necessarily mean the region is devoid of repetitive sequence. When comparing GATK-SV, which was used to create gnomAD-SV, almost all deletions (98%+) were imperfect matches, regardless of sequence context and database. Consistent with benchmarking against the HG002 truth set (Figure 3A), the Manta and priority_merge callsets perfectly matched over 80% of common HG002 deletions that were found in CoLoRS and 1KGP-ONT. In contrast to deletions, at least 20% of common insertions in the HG002 GATK-SV callset were perfect matches in all three databases (Figure 4B). The insertions output by GATK-SV are slightly more concordant with gnomAD-SV than the long-read sequencing databases. On the other hand, insertions from Manta and priority_merge have substantially fewer perfect matches to gnomAD-SV than 1KGP-ONT and CoLoRS. A small increase in perfect matches to CoLoRS and 1KGP-ONT is also observed for both deletions and insertions in the priority_merge callset compared to Manta. Surprisingly, there were only minor differences in matching accuracy between the Unique, and SimpleRepeats or SegmentalDups annotated regions for the Manta and priority_merge insertions and deletions in the long-read sequencing databases. While CoLoRS and 1KGP-ONT both show good agreement between perfect matches for Manta and priority_merge, insertions consistently have a greater proportion of perfect matches to CoLoRS than 1KGP-ONT. This indicates representation differences exist between long-read sequencing databases for common insertions that appear to have minimal, if any, dependence on repetitive sequence context.

**Figure 4.**
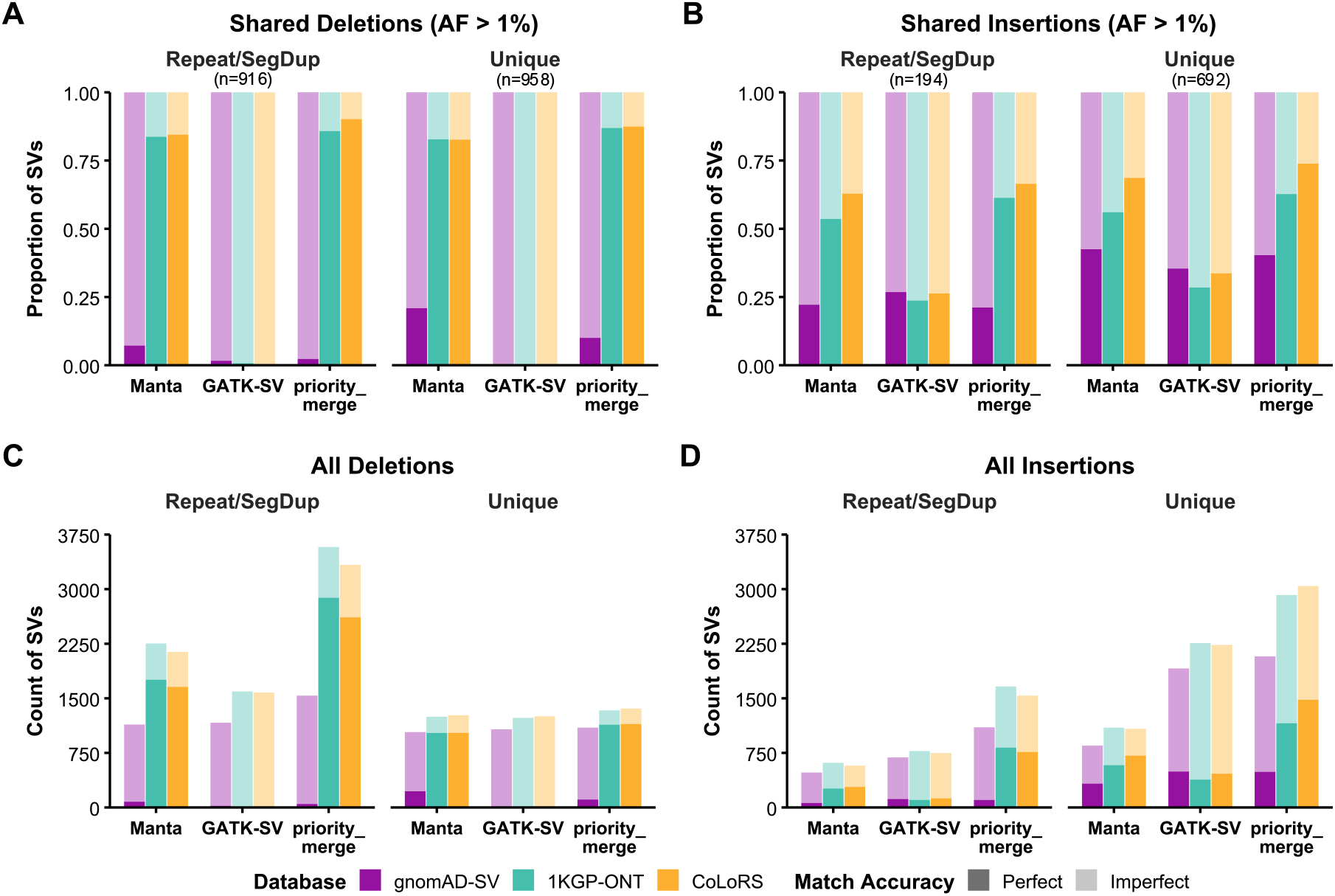
Matching accuracy and ‘missingness’ of Manta, GATK-SV and priority_merge insertions and deletions compared to the gnomAD-SV, 1KGP-ONT and CoLoRS databases. Shared SVs in **A**) and **B**) are determined by intersecting the three callset-specific representations of each HG002 truth set SV with their database matches. The three callset representations must have matched to the same SV VCF record in each database (Methods). **A)** Differences in matching accuracy between common HG002 deletions. **B)** Differences in matching accuracy between common HG002 insertions. **C)** Differences in matching accuracy between all HG002 deletions. **D)** Differences in matching accuracy between all HG002 insertions. Repeat/Segdup annotations represent the HG002 truth set SV coordinates that had at least one bp of overlap with UCSC SimpleRepeats or Segmental Dups tracks. Imperfect match accuracy SVs are any callset representation that did not perfectly match the database SV but matched at the permissive matching threshold (70% size and 500bp reference distance (insertions) or 70% reciprocal overlap (deletions)). The 1KGP-ONT and CoLoRS databases are derived from nanopore and PacBio HiFi long-read sequencing of 500 and 1,427 samples, respectively.

Identifying novel disease-associated genes and pathogenic variants in known disease genes requires comprehensive variant detection and estimation of allele frequencies to interpret the genetic variation discovered. Thus, we next matched the three callset representations of all HG002 truth SVs detected in the three callsets to the databases to evaluate ‘missingness’ in addition to matching accuracy. All values in this section that refer to long-read sequencing databases indicate averages (mean) across 1KGP-ONT and CoLoRS unless stated otherwise. Within repetitive genomic regions, Manta perfectly matched more deletions (1,707) in the long-read sequencing databases than the total number of deletions matched by GATK-SV (perfect: 6; imperfect: 1,587). Manta also perfectly matched more 1KGP-ONT or CoLoRS deletions than the total matched by any callset to gnomAD-SV (max: priority_merge - 1,537) (**Figure 4C)**. The priority_merge callset likewise contained more perfect matches (2,748) than the total number of deletions matched by Manta (2,196) to the long-read databases. Relative to repetitive regions, there were marginal differences in the total number of deletions matched in unique regions between the three callsets to the SV databases. Yet, in all three callsets, the Unique regions still had 20.0% more deletion matches on average to 1KGP-ONT and CoLoRS than gnomAD-SV. The priority_merge callset had more perfect matches to the long-read databases than total matches to gnomAD-SV, including higher total matches than GATK-SV (+8.5%) or Manta (+7.2%). For insertions in repetitive sequence contexts, there were 60.3% (414) more matches to gnomAD-SV in the priority_merge callset than GATK-SV (Figure 4D). This suggests many insertions within these regions were not rediscovered by GATK-SV in Koenig et al. [29] or were incorrectly joint-genotyped for the HG002 sample, but are present in gnomAD-SV and may have unreliable allele frequencies. Moreover, in repetitive regions, priority_merge had a greater count of insertions that perfectly matched the long-read databases (793; 49.6% of total) than the total number of matches by GATK-SV (max: 775) or Manta (max: 614) to any database. Although Manta found fewer total insertion matches in each database than GATK-SV, it produced substantially more perfect matches to 1KGP-ONT and CoLoRS (Repeat/SegDup: 271, +135.7%; Unique: 648, +52.8%). The priority_merge callset, however, found roughly twice as many perfect insertion matches to the long-read databases compared to Manta. Furthermore, GATK-SV had 735 (-24.6%) fewer total insertions in unique regions than priority_merge that were matched to 1KGP-ONT and CoLoRS. It should also be noted there are 44, 134 and 356 SVs found in the false positive VCFs of the GATK-SV, Manta and priority_merge callsets, respectively, that are true positives in the HG002 truth set but did not match to any database.

## Discussion

Here we demonstrated a proof-of-concept intra-sample SV merging strategy using highly accurate individual callers (gcSV, Manta, SurVIndel2, INSurVeyor) that prioritises selecting a representative SV from redundant matches based on benchmarking performance against multiple matching thresholds. This method improves upon current state-of-the-art ensemble pipelines, increasing the comprehensiveness of SV callsets without sacrificing matching accuracy relative to the best performing tools in the ensemble. The HG002 ensemble callset we generated (priority_merge) matched more SVs in two long-read sequencing databases (CoLoRS, 1KGP-ONT) and had better agreement on their breakpoints and sizes when compared to a short-read database (gnomAD-SV) and its respective ensemble pipeline (GATK-SV) callset for the same sample. Most of the improvements in deletion recall, including perfect matches, occurred within repetitive regions of the genome. Insertion improvements were more evenly distributed across both unique and repetitive regions. We also found that the Manta and priority_merge callsets are more concordant with 1KGP-ONT and CoLoRS than gnomAD-SV, highlighting the additional value of long-read sequencing databases for obtaining allele frequency estimates and SV interpretation.

Although using newer callers and priority merging SVs based on benchmarking against multiple matching thresholds improves upon existing ensemble methods, this study has several limitations. First, an intra-sample merging strategy prioritising recall inherently increases false positives when using multiple SV callers. Like other ensembles, any implementation of this strategy will need to carefully select which SV callers to include and determine an appropriate balance for this trade-off. As the availability of high-quality, comprehensive SV truth sets grows, machine learning models are increasingly promising candidates for reducing false positives here. These models can also potentially be extended to integrate sophisticated merging approaches that consider genome annotations [30,31], caller-specific VCF INFO fields like breakpoint coverage [32], and other features that may better predict breakpoint and size accuracy between redundant SVs [33]. Second, we did not evaluate genotyping accuracy because some tools do not output genotypes (notably INSurVeyor and SurVIndel2), necessitating a secondary re-genotyping step. Third, inversions were excluded from benchmarking because they are absent in the HG002 truth set. Inversions may also require more focused benchmark development and evaluation efforts, particularly for complex events [34]. Fourth, we did not explicitly benchmark copy number variants (CNVs). CNVs require benchmark datasets and long-read sequencing databases representing CNVs with copy numbers, including tools that output and assess copy numbers. Moreover, as long-read sequencing has matured, a conflict between definitions of variant type and biologically equivalent representations of CNVs is becoming apparent. This partially stems from the sequence resolution improvements with long-read sequencing that largely occur in tandem repeats (TR) [35] and low-complexity regions [31]. For instance, copy number differences of TRs can be output as duplications with short reads, but will tend to be represented as sequence-resolved insertions by long-reads (like the SV truth sets used here). This difference in sequence resolution and variant type also occurs between short-read methods for the same SV [7,32]. Fifth, we did not assess sequence accuracy, which shares some issues, among others, related to the CNV limitations. Finally, the effect of inter-sample merging on SV breakpoints and sizes was not analysed here.

There are three main steps where SV breakpoint and size inaccuracy errors can be made with ensemble methods: 1) choice of SV callers, 2) intra-sample merging (identifying the same SV across multiple callers), and 3) inter-sample merging (identifying the same SV across multiple samples). The performance of Parliament2 and MOPline on a single sample compared to their individual tools, particularly Manta, strongly indicates their inaccuracy is due to intra-sample merging. However, the GATK-SV callsets we used for HG002 [27], and NA12878 / NA12877 [5] were obtained from previous studies involving many samples. They were extracted from joint-genotyped VCFs that have undergone inter-sample merging, which could have introduced another error source for its SV breakpoints and sizes. From the analysis conducted here, inter-sample merging does not appear to be a satisfactory explanation for the difference in matching accuracy between the GATK-SV callsets and gnomAD-SV. It is expected that highly similar TR alleles will be collapsed across samples and produce incorrect SV breakpoints and sizes at some loci for an individual sample [24]. If inter-sample merging was the source of the errors, we would expect to find comparable matching accuracy between GATK-SV/gnomAD-SV and Manta with the long-read sequencing databases in Unique regions and significant differences in Repeat/SegDup regions. We do not observe this for the HG002 sample. Furthermore, GATK-SV recall in Figure 1 is consistent with previous observations from the HGSVC, which found single sample breakpoints output by GATK-SV had better agreement with svtools (a short-read ensemble pipeline) than PacBio assemblies [5], supporting intra-sample merging as the cause. The Manta and priority_merge matching accuracy of common HG002 insertions and deletions in 1KGP-ONT and CoLoRS (**Figure 4A & Figure 4B**) also suggests the simplest inter-sample merging strategy (median or mode of breakpoints/sizes) would have converged on the ‘true’ representation for most common SVs.

As studies of rare diseases and complex traits begin to integrate and jointly analyse all types of genetic variation (SNVs, small indels, TRs, SVs, CNVs) [7,36], the overlap between SV discovery and TR genotyping tools presents a significant variant merging challenge, particularly for inter-sample merging. A distribution of repeat sizes and sequence differences at TR loci is required to identify TR expansion thresholds in rare diseases, and length and sequence associations in complex traits. SV discovery tools inherently detect expansions and contractions of TRs, but information currently output in VCFs by these tools make genome-wide inter-sample merging and analysis extremely difficult without substantial postprocessing efforts. For example, when merging across samples, applying permissive matching thresholds collapses true differences in repeat sizes between TR alleles into a single SV with an inflated allele frequency. Additionally, as sample sizes of short- and long-read sequencing population databases continue to increase, potential single base differences between common and rare SVs within unique and repetitive regions will be lost. Surprisingly, we observe high agreement (50%+) on exact SV breakpoints and sizes (particularly deletions – 80%+) in repetitive regions between the HG002 truth SVs, the two long-read sequencing databases, and the Manta and priority_merge callsets. It is unclear how much of this is explained by biallelic sites, low polymorphism rates in the population, and inter-sample merging strategies for the 1KGP-ONT and CoLoRS samples, which may have selected the breakpoints and size of the most common TR allele that is also shared by the HG002 sample. Due to our annotation method (Methods), a small proportion of these will also be segmental duplications. Not all Repeat/SegDup and Unique annotated SVs will be correctly annotated as TRs either (TR definitions can vary dramatically) [37]. Developing robust inter-sample merging strategies will be necessary to eliminate redundant variant representations, retain allelic diversity and prevent allele frequency inflation [24]. Like intra-sample merging, machine learning models could also be useful for determining when highly similar SVs between samples should be merged into a single record and what the breakpoints and size should be, or whether they should be left unmerged. In-depth analysis of the respective roles for short-read SV discovery and TR genotyping tools, and how to handle the complexities of their overlap, are needed for future joint-analysis studies of SVs and TRs with SNVs and small indels.

Nevertheless, assuming similar read processing, we expect the SV database analysis and intra-sample merging strategy we present here to be broadly generalisable given the high concordance in matching accuracy between the HG002 truth set, the Manta and priority_merge callsets, and both long-read sequencing databases. This is further implied by the similarity in matching accuracy between the HG002 T2T-Q100 and Platinum Pedigree truth sets and the callset benchmarking. We recommend applying multiple matching thresholds (or finer breakpoint and size resolution analyses) when benchmarking SV calling tools. By adopting this recommendation, the choice of structural variant callers applied individually or within ensemble pipelines to Illumina short-read WGS data becomes dramatically simplified. Any new tool for SV discovery should demonstrate its perfect matching threshold performance or its utility should be justified by showing significant unique contributions to callset comprehensiveness at lower thresholds relative to more precise tools. Furthermore, breakpoint and size resolution analyses, like those presented here, should become standard when developing ensemble pipelines and generating multisample SV callsets to validate intra- and inter-sample merging strategies. Specifically, comparisons should include benchmark samples and examine the SV counts and shift in breakpoints and sizes between the merged and unmerged benchmark samples. Ultimately, prioritising exact base-pair resolution will improve comparisons between short-read WGS studies, ensure accurate functional annotations, and help facilitate the integration of diverse variant classes in future disease and trait studies.

## Data Availability

All unprocessed VCFs, processed truth sets, and BED files used for benchmarking and analysis are available on Zenodo (DOI: https://doi.org/10.5281/zenodo.21126342). The custom Python scripts used for VCF standardisation, caller priority merging, and database intersections are available at GitHub (https://github.com/jcslimmer/prioritySV) and archived on Zenodo (DOI: https://doi.org/10.5281/zenodo.21126342).

## Author contributions

J.C.S.: Conceptualisation, Methodology, Software, Formal analysis, Investigation, Data curation, Visualisation, Writing – original draft. J.L.J.: Supervision, Writing – reviewing & editing. W.R.R.: Supervision, Writing – reviewing & editing. K.P.B.: Supervision, Funding acquisition, Writing – reviewing & editing. N.B.B: Supervision, Funding acquisition, Visualisation, Project administration, Writing – reviewing & editing.

## Acknowledgements

The manuscript text, figures and code were prepared with the assistance of Gemini models from Google DeepMind. AI was used to refine language, improve clarity, enhance readability and proofread errors in accordance with the authors’ instructions. AI assistance was similarly used to draft and debug code associated with the analyses and figures generated for this manuscript. All content, scientific claims, and conclusions have been reviewed and verified by the authors to ensure accuracy and originality.

---

**Supplementary Table 1.**
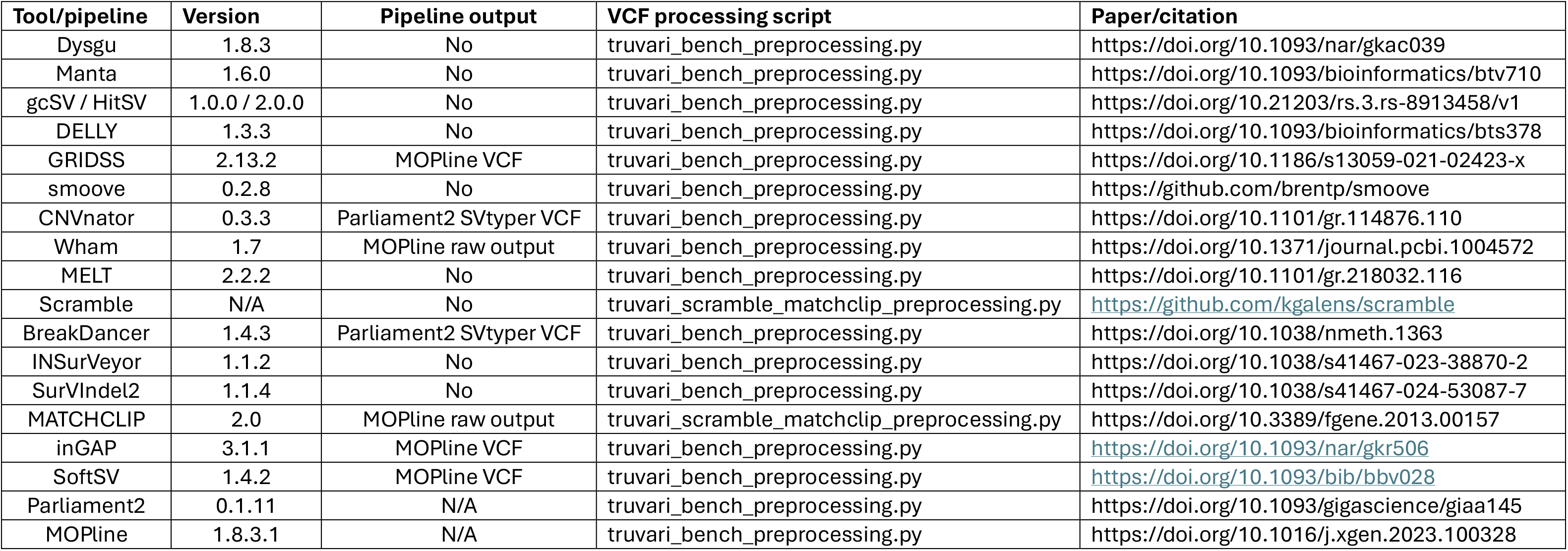
SV caller and pipeline version details. The pipeline output column describes which VCF was used for benchmarking when using one of the ensemble pipelines to run the SV caller. N/A – not applicable.

**Supplementary Figure 1.**
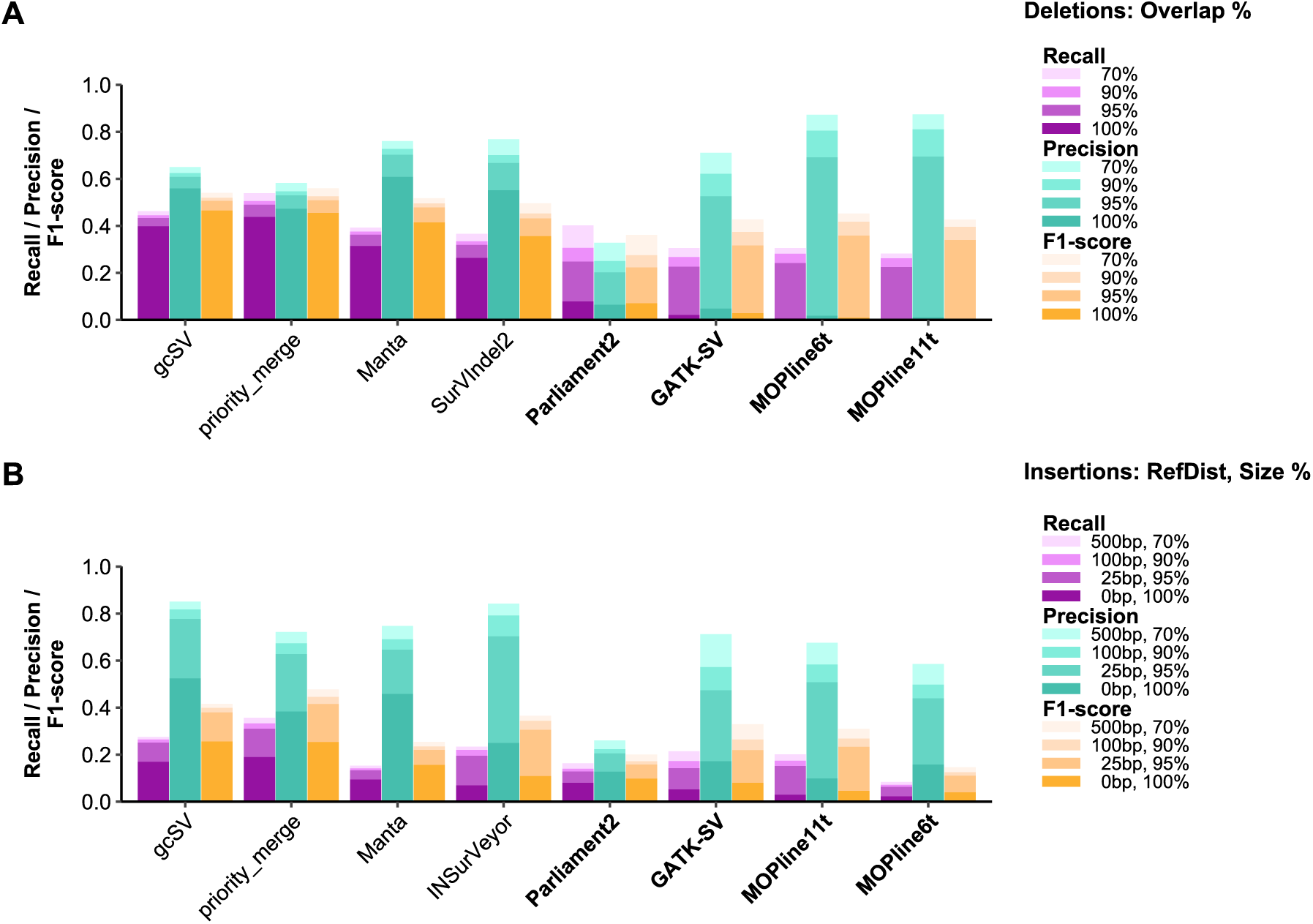
Recall, precision and F1-scores on the NA12878 sample truth set when priority merging insertions and deletions output by multiple SV callers. A ‘priority_merge’ callset was constructed by splitting individual caller VCFs by SV type and size bin, followed by merging them with Truvari using a caller priority order (e.g. 50-500bp deletions: gcSV > Manta > SurVIndel2; 50-200bp insertions: gcSV, Manta, INSurVeyor) informed by their benchmarking performance. The caller priority orders for the priority_merge callsets can be found in the **Methods** section. Colours indicate the recall, F1-score and precision for each callset along the x-axis. From lightest to darkest, the shade of each colour represents increasingly stringent matching thresholds. Callsets for **A)** deletions and **B)** insertions are ordered by descending F1-score at the perfect match threshold. Ensemble pipelines that integrate multiple SV calling tools are highlighted in bold (all include Manta).

**Supplementary Figure 2.**
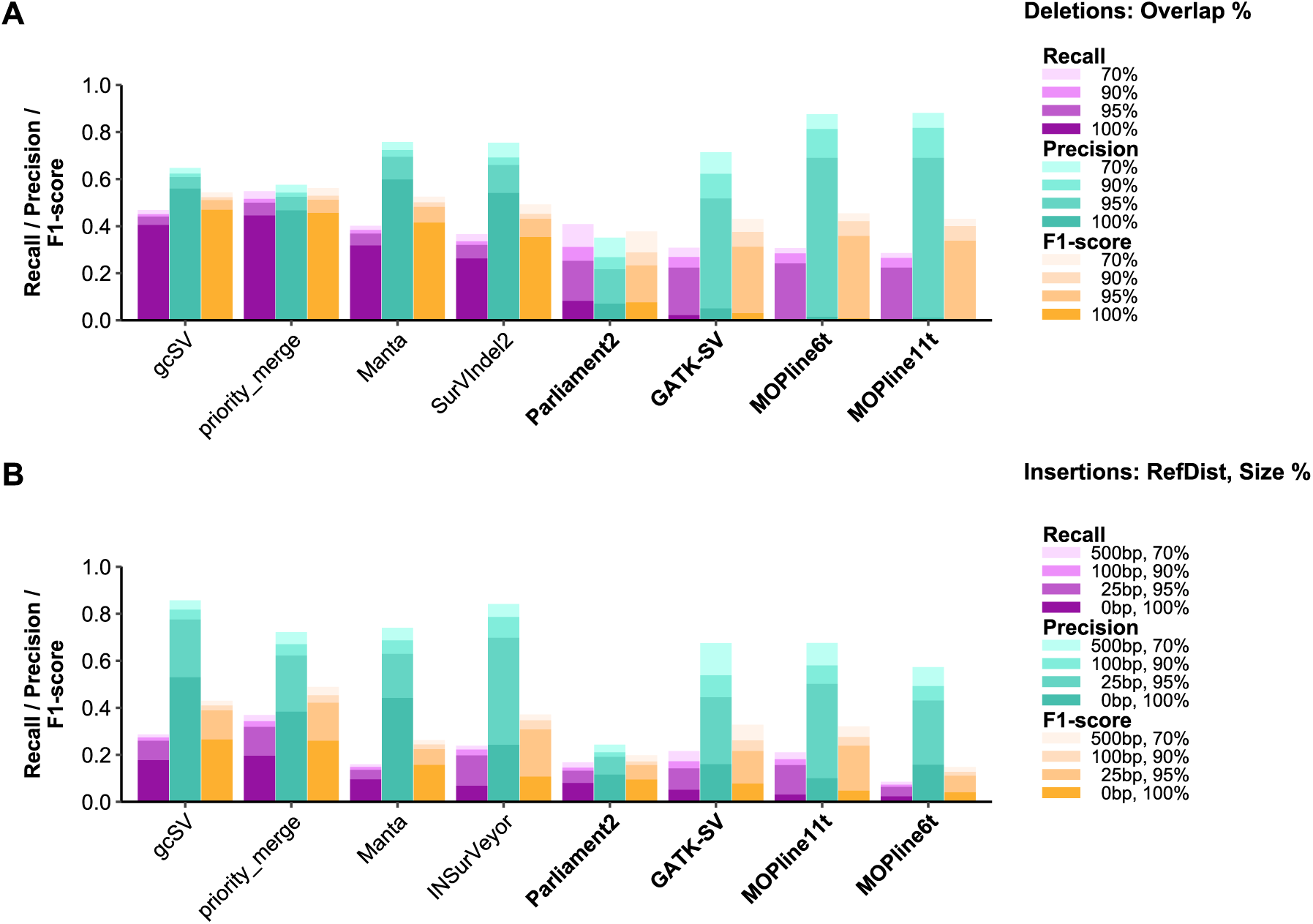
Recall, precision and F1-scores on the NA12877 sample truth set when priority merging insertions and deletions output by multiple SV callers. A ‘priority_merge’ callset was constructed by splitting individual caller VCFs by SV type and size bin, followed by merging them with Truvari using a caller priority order (e.g. 50-500bp deletions: gcSV > Manta > SurVIndel2; 50-200bp insertions: gcSV, Manta, INSurVeyor) informed by their benchmarking performance. The caller priority orders for the priority_merge callsets can be found in the **Methods** section. Colours indicate the recall, F1-score and precision for each callset along the x-axis. From lightest to darkest, the shade of each colour represents increasingly stringent matching thresholds. Callsets for **A)** deletions and **B)** insertions are ordered by descending F1-score at the perfect match threshold. Ensemble pipelines that integrate multiple SV calling tools are highlighted in bold (all include Manta).

